## Supplemental text for "TWINKLE: An open-source two-photon microscope for teaching and research"

### Supplemental Online Material

This supplemental material to the article "TWINKLE: An open-source two-photon microscope for teaching and research" describes the assembly, function and alignment of all parts of the microscope. Technical specifications are provided through the CAD files and the Zemax simulations in the online repository at
https://github.com/BrainCOGS/Microscope.

**6 Recommendations for use** **28**

**7 Deriving the relationship between intensity and dispersion** **29**

### 1 Preface

#### 1.1 Safety

Class 4 lasers are extremely dangerous. A reflection on a surface can be enough to inflict permanent eye damage. Safety protocols must be observed. Consult with your institution's environmental health and safety officer and take proper training before performing any work with a laser system. Learn from people experienced with these devices and work carefully.
During alignment, always wear goggles (ours were ML1 by NoIR LaserShields with an optical density $> 7$ in the relevant range around 920 nm), remove jewelry and watches, only use-reflective tools, do not store unnecessary items on the optical table, and use the low laser power ("alignment mode") whenever possible. Once the assembly progresses further, full power mode is necessary. Extra care must be taken then. Always use Laser Viewing Cards to track the laser. We use Thorlabs VRC4. Always be conscious where the beam is, and where the beam goes. The beam must be terminated in beam blockers and should be shielded whenever possible with opaque plastic or anodized aluminium tubes to increase safety.

#### 1.2 Miscellaneous tools and tips

The following tools are used throughout the build process:

- A high quality duster to remove any residue from optical surfaces. We use a Miller-Stephenson Aero-Duster Ultra MS-222L, in addition to lens tissues (MC-5, Thorlabs) and Methanol (Sigma-Aldrich).
- To glue optical filters we used an extra fast setting epoxy (Hardman Double/Bubble Epoxy). To seal small gaps, we use black silicone putty (V-tie self-setting silicone rubber).
- We used either McMaster or Thorlabs screwdrivers and hex keys.
- For alignment, we used laser viewing cards (VRC4, Thorlabs), cage system alignment plates (VRC4CPT, Thorlabs), and irises (SM1 Graduated Ring-Actuated Iris Diaphragm, SM1D12C, Thorlabs).

The following general tips might be of help as well:

- Shield the beam as soon as alignment of a particular section is done. This increases safety, and avoids accidentally bumping into aligned components.
- *Walking the beam*: Any beam can be aligned through two irises using two mirrors. The first mirror the beam meets is sometimes called the "position" mirror as it affects primarily the position of the beam. The second "angle" mirror instead primarily affects the angle. Assigning each mirror a separate function (position versus angle) and by adjusting them iteratively, is a good rule of thumb for driving the beam through the 2 irises and eventually hitting a specific point at the desired angle [1].
- Be mindful of reflections from all surfaces. Put back-reflecting component at a slight angle to the beam (e.g. the half wave plate) to avoid back-reflections directly into the laser. The back-reflection can then be caught with an iris placed in front of the laser.
- Make sure the beam is at the same height across the table. This can be done with an iris and an IR card.

| Item | Info | Cost |
|---|---|---|
| 920 nm fs-pulsed Laser | Spark Alcor 920 + Warranty | 90 k |
| ScanImage DAQ-System & optimized PC | MBF Bioscience | 20 k |
| Piezo collar | Physik Instrumente | 16 k |
| 2× PMTs + preamplifiers | Hamamatsu & Thorlabs | 12 k |
| Pockels cell | Conoptics | 11 k |
| Objective | Nikon | 8 k |
| Machining custom parts | Machine Shop; as per CAD files | 7 k |
| Optical filters | AVR optics & Semrock | 7 k |
| Lenses + optomechanics | Thorlabs & Newport | 7 k |
| Galvo-Resonant Scanner set | Novanta | 6 k |
| Electronics/cables/tools | Newark & McMaster | 5 k |
| Mounting rack | Emcor Enclosures | 3 k |
| Laser shutter | Uniblitz/Vincent | 2 k |
| Laser safety goggles | Edmund Optics | 1 k |
| Laser power meter | Coherent | 1 k |
| Agarose & fluorescent beads | Thermofisher | 1 k |
| **Total** | | **197 k** |

**Table 1. Overview of the approximate cost in US-$ (2024).** Note that this list excludes the optical table, and any other infrastructure costs such as compressors for air, and associated power supplies. This is just a summary. The detailed bill of materials is provided in the git repository.

- Take extra care for directing the beam through the center of all lenses and active optical components.

##### 1.3 Parts and bill of materials

All custom metal pieces are parts of the CAD assembly of the microscope. Ours were manufactured from aluminium stock in the university machine shop. Aluminium has the added benefit of anodization, which can help control scattered light. An overview of the microscope's cost is shown in Tab. 1. Keep in mind that these prices can significantly change over time, while some products can become unobtainable and need to be replaced with a different or newer model.

##### 1.4 Structure and use of this document

In the following sections, we will cover the setup and alignment of the microscope. This is split into three parts. The first part covers all components on the optical table that serve to condition the beam. The second part focuses on the 95 mm vertical post that holds the scanning mirror and scan lens assemblies. The third part is the microscope head with the attached collection optics. We will begin with the assembly on the optical table. All along, we assume the reader will have downloaded and opened the CAD model as we will rely heavily on it. Further, check carefully the instruction manuals of the scanning mirrors, photomultiplier tubes (PMTs), power supplies, Pockels cell, ScanImage, the Piezoelectric objective collar, and the filter mounting instructions. Before building your own microscope, read these instructions fully. Check the CAD files, and make sure that the instructions make sense. This document follows closely our own alignment procedure, and we have done our best to be accurate. However, use common sense. Do not follow these instructions blindly (or you might follow these instructions blindly in the future). Like all open instrumentation (under CC-BY-4.0), this work is

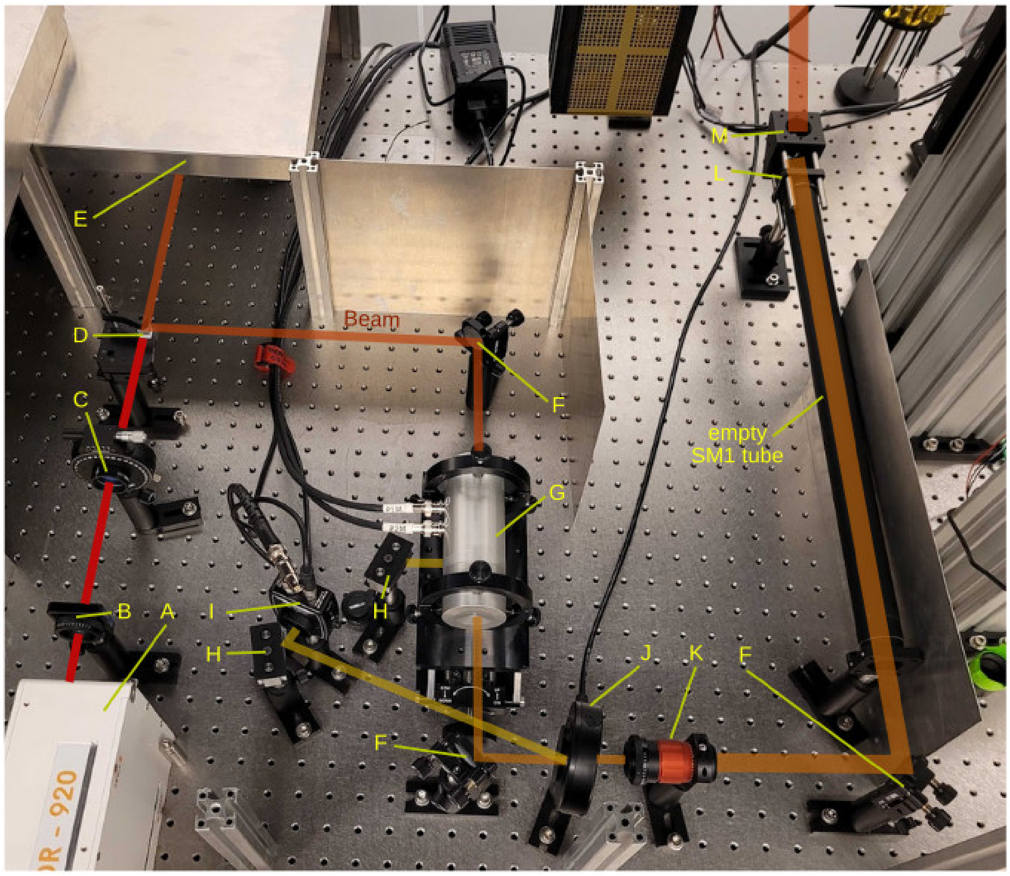

**Figure 1. Overall view of the beam conditioning path on the optical table.** Shown is a photograph of the components used to condition the beam for our microscope. Key components of the system are labeled with capital letters. The laser is shown as a red transparent line with intensity color coded. The beamsplitter cube (component D) splits the beam into two, halving its power. The Pockels cell (component G) reduces the power further. The reflections out of the Pockels cell and from the shutter (item J) are shown in yellow. The telescope (item K) magnifies the beam $\times 2$. The empty SM1 tube is an anodized aluminum SM1 lens tube cover (SC600, Thorlabs). All parts are explained in the text together with a note on their alignment.

distributed in the hope that it will be useful, but WITHOUT ANY WARRANTY, to the extent permitted by law; without even the implied warranty of MERCHANTABILITY or FITNESS FOR A PARTICULAR PURPOSE.

### 2 Assembly on the optical table

#### 2.1 Beam conditioning

An overview of the system for beam conditioning is shown in Fig. 1 with key items labeled by capital letters. The following lines will cover the items, and their alignment.

**A)** Laser head emitting the laser beam (red).

**B)** Iris (SM1 Graduated Ring-Actuated Iris Diaphragm, SM1D12C, Thorlabs) inside a SM1-Threaded 30 mm cage plate (CP33, Thorlabs).

**C)** ⌀1 inch mounted achromatic half-wave plate with SM1-threaded mount (AHWP10M-980, Thorlabs) inside a high-precision rotation mount (PRM1, Thorlabs). First, use an iris to check that the beam travels centrally through the half-wave plate. Then place the half-wave plate at a slight angle to make sure that the beam reflected of the surface of the half-wave plate does not travel back into the laser. Confirm by using a detector card to find the (weak) back reflection. While doing this, terminate the beam traveling through the half-wave plate with a beam block.

**D)** 10 mm polarizing beamsplitter cube (PBS103, Thorlabs) on a kinematic platform mount (KM100B, Thorlabs) held in place with a large adjustable clamping arm (PM4, Thorlabs). Arrows on the cube indicate which way the beam should go in and where the split beams travel. Similar to the half-wave plate, confirm that the reflection does not travel back to the laser using a detector card. Then adjust the kinematic mount to make sure that the transmitted and split beams travel parallel to the optical table. This is best checked with an iris far after the cube, and a detector card. Finally, adjust the angle on the half-wave plate such that transmitted and split beams have equal power. This has to be done at full-power mode of the laser. We used a Coherent FieldMaxII with a PM10 Thermopile power sensor.

**E)** Beam block (hidden under the aluminium shield). This blocks the second beam which is not in use in our system (LB1, Thorlabs).

**F)** ⌀1 inch protected silver mirror (PF10-03-P01, Thorlabs) inside a kinematic mirror mount (KM100, Thorlabs): Reflects the beam.

**G)** Pockels cell (350-80-LA-02 with 302RM driver, Conoptics) sitting on a holder (M102A, Conoptics). Properties of light travelling through this device are controlled electrically through the two connected wires. The alignment of the Pockels cell requires two steps: The Pockels cell has to be aligned with the optical axis of the laser, and be rotated relative to the laser's polarization such that the dynamic range is maximized [2,3]. The first step can be done with an alignment tool of identical geometry to the Pockels cell (Conoptics modulator alignment tool 103). The second step requires the Pockels cell to be connected, and a power measurement performed. Importantly, these steps have to be done under full power because the beam profile in alignment mode might be slightly different from the full beam.

- First, install the holder on the table, and position on it the Conoptics Pockels cell alignment tool (Conoptics modulator alignment tool 103).
- Use a power meter on the output side, maximize power going through the central bore by manipulating the 4 bottom screws on the holder.
- Then switch the laser to full power and optimize the position of the screws for maximum transmission.
- Tighten the screws around the alignment tool, and mark on the screws the optical position. This will be important later to reproduce the exact same position of the alignment tool with the Pockels cell.
- Connect the Pockels cell (M350-80-LA, Conoptics) to the driver as per the Conoptics instructions (`p1m` and `p2m` go to `J1m` and `J2m`, `p1` and `p2` go to the controller). The controller settings are shown in Fig. 2.
- Next, determine the best orientation of the Pockels cell: For several rotations of the Pockels cell, measure its transmission while turning the dial on the Conoptics controller from one end to the other. Iterate this for different orientations of the Pockels cell until an optimum is found at which the dynamic range is greatest. This optimal orientation allows to adjust the laser power in the widest range. An

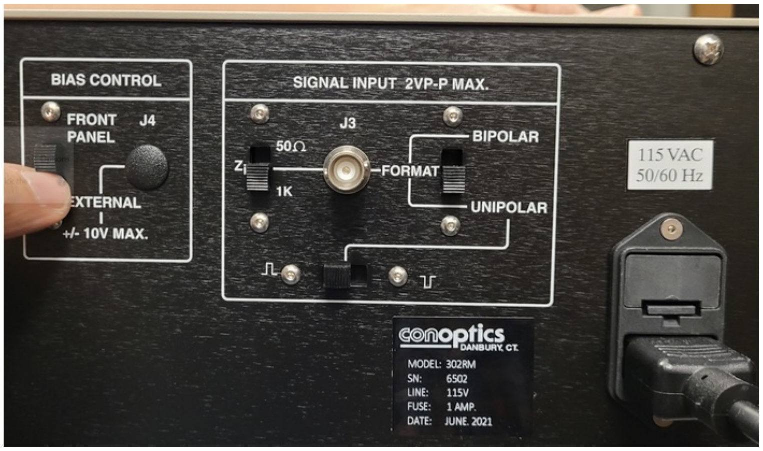

**Figure 2. Settings on the Pockels Cell controller.** Shown are the controls of the Pockels Cell driver as set in our system. For reference, see [2]. We recommend for people building a microscope to take many such pictures of all settings and elements.

example of different transmission curves is shown in the main manuscript, Fig. 9A.

- The Pockels Cell splits the input beam into a transmitted and a rejected beam. The transmitted beam is used, while the energy of the rejected beam is dissipated in a beam block. By changing the control voltage of the Pockels Cell, the power ratio between the transmitted and rejected beam can be adjusted. The rejected beam leaves the Pockels cell through a bore at its side. During alignment, make sure that the rejected beam is properly terminated with a beam block.
- To validate the alignment, compute the extinction ratio of the Pockels cell, i.e. the ratio of maximal and minimal light transmission, and compare that number with the data sheet provided by Conoptics. Ours operate in a regime of $\approx 450$.
- Next, set the Pockels cell controller to the control voltage corresponding to minimum light transmission.
- Tighten the holder, and confirm that the Pockels cell is still well-aligned.

**H)** Two beam blocks (LB1, Thorlabs). Confirm that the right beam block terminates the rejected beam leaving the Pockels cell. The left beam block is used for calibrating the Pockels cell in ScanImage (the control software used by our microscope [4, 5]), together with a light sensor (I).

**I)** Si Switchable Gain Detector, 350 - 1100 nm (PDA36A2, Thorlabs). This is used for the laser power calibration of the light transmitted through the Pockels cell. The detector is facing the beam block (H) and monitors the light that comes from the shutter back reflection when closed. Make sure that the gain on the detector is chosen reasonably for laser power and ambient light level. In our system, a gain of 50 dB works well. If the ambient room light is too bright, a SM1-mounted infrared long-pass filter in front of the photodiode can help (e.g. FELH0850, Thorlabs). The goal of setting the gain is well-defined sigmoidal control curve in ScanImage: Light transmission through the Pockels cell starts at a minimum, and increases as the Pockels control voltage increases.

**J)** 6 mm Shutter (LS6S2ZM1, Vincent Associates) connected to driver (VCM-D1, Vincent Associates) for control of the shutter during imaging. To check that the shutter is properly aligned leave it open and make sure that the beam is not clipped. When closed, the back-reflection is sent to a beam block. For this reason the shutter is not mounted exactly perpendicular to the incident beam but at a slight angle.

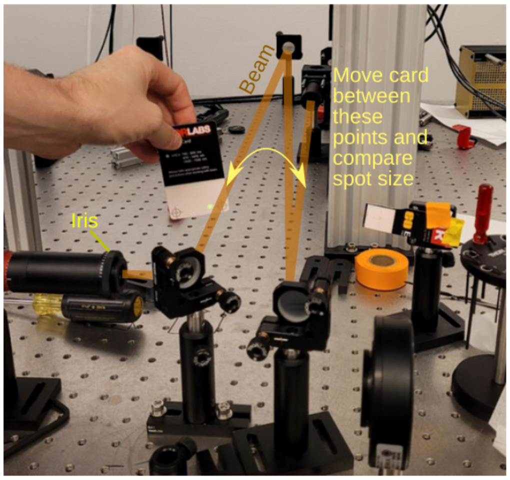

**Figure 3. Method for making sure the beam is collimated.** Using a set of mirrors, the beam (yellow line) can be reflected back and fourth to increase the distance travelled. Along this path (in this example of $\approx 2.5$ m), the beam must not change size ($< 1$ mm). This is validated with an IR detection card and switching between the forward and return path of the beam. This method allows to control beam divergence after the 2× telescope to below 1 mrad.

**K)** 2× Beam expander (2× Achromatic Galilean Beam Expander, GBE02-B, Thorlabs, inside SM1RC, Thorlabs). Aligning the beam expander requires two steps. First, it has to be aligned on the beam axis. Second, the beam needs to remain collimated after the beam expander, which is achieved by tuning the expander collar. For the first alignment, we use two irises, one screwed in the front, and one (on a reasonably long SM1 extension tube) behind the expander. The beam expander has to be positioned such that the beam travels through both iris, and such that the irises cut into the beam symmetrically. Next, we need to adjust the collimation of the beam using the expander collimation collar. It needs to be tuned so that the beam is the same size at the exit and after a long distance using several mirrors, see Fig. 3.

**L)** Iris (SM1D12C mounted on a CP08 plate, both Thorlabs) used to check alignment.

**M)** Mirror (PF10-03-P01, Thorlabs, inside KCB1) to reflect the beam up at a right angle. For the next steps (in particular finding the optimal scanning mirror position), it is important that the beam travels up perpendicularly to the optical table, and parallel to the 95 mm optical rail column that holds the scanning mirrors (next section).

When finished aligning the beam through all the optical components listed above, clamp them down on the table securely. Then shield the beam path with additional SM1 and SM2 tubing, and tubing cover (e.g. Thorlabs SC600, SM1 Anodized Aluminum Lens Tube Cover) to increase safety and to protect the aligned optical components.

#### 2.2 Assembly of the scanning post

##### 2.2.1 Components

Following beam conditioning on the optical table, the beam is reflected up from the optical table towards the ceiling into a box housing the scanning mirrors. This is shown in Fig. 4A,B. Following the scanning mirrors, the beam travels up through the scan lens, and directed to the microscope head with another mirror.

**A)** Mirror (PF10-03-P01, inside KCB1, with CP12 attached) - turn the beam at a right angle.

**B)** Broadband filter (WG11010-B, Thorlabs, inside SM1L20 and SM1S20) - filter to keep dust out of the box housing the scanning mirrors.

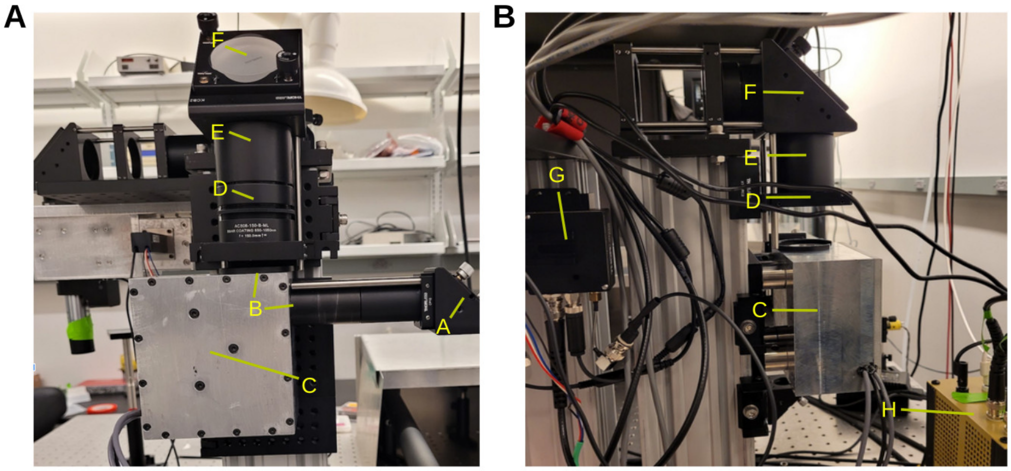

**Figure 4. Overall view of the scanning post. A)** Side few. Key components of the system are labeled with capital letters. These are explained in the text together with a note on how to align these components. Importantly, the machined block (C) houses the scanning mirrors, and the scan lens assembly is placed in the SM2 house labeled (E). **B)** Frontal view. The PMT preamplifiers (component G) and the scanning mirror power supply (component H) are visible.

**C)** An aluminium box containing the resonant and slow galvanometer set (CRS8K/6215H Resonant-Galvo Set, Novanta), mounted to 95 mm rail (XT95RC4, Thorlabs). A detailed description is provided below.

**D)** Empty tubing (SM2V05, adjustable SM2 Lens Tube, Thorlabs).

**E)** Scan lens - See detailed description below.

**F)** Mirror in its housing (PF20-03-P01 and KCB2, both Thorlabs). While we used a round ⌀2 inch mirror, this should be replaced with an elliptical mirror and housing (PFE20-P01 and KCB2EC, both Thorlabs). This would increase the field-of-view because the cross-section of an ellipse at 45 degrees is a circle. At low magnification, a round mirror will lead to vignetting.

**G)** PMT preamplifier (TIA60, Thorlabs) attached to a 95 mm rail.

**H)** Power supply for the scanning mirrors (part of the Novanta scanning mirror set) housed in an electronics prototype box.

##### 2.2.2 Alignment

- First, mount the 95 mm rail to the table. Then attach a bracket (all from the XT95 series, Thorlabs) to the side to hold the scan lens (component D in Fig. 3), and one on top with a cage system to hold a mirror (component F in Fig. 3). For alignment with cage systems, we use a cage system alignment plate (VRC4CPT, Thorlabs). The rest of the assembly here will be done without any lenses. The goal is a well-aligned beam that travels through the center of all parts, and parallel to the optomechanical surfaces.
- The box housing the scanning mirrors is connected, trough SM1 tubing, to a mirror (component A in Fig. 3). To align this upper periscope mirror with the lower periscope mirror, mounted on the table, connect a piece of SM1 tubing to

the upper mirror facing down, and a second piece of SM1 tubing to the table mirror facing up (component M in Fig. 1). These two pieces meet in mid-air. The gap in between can be reduced to zero using adjustable length SM1 lens tube (e.g. SM1V10, Thorlabs, not shown in CAD) and can be further closed by a small piece of lens tube cover (SC600, Thorlabs).

- Next, we adjust the mirror (component A in Fig. 4A) so that light travels through the center of the scanning mirrors and up the 95 mm rail. This is done with irises. One is attached at the entry port of the scanning mirror box, e.g. in the SM1 tube (component B in Fig. 4A). Another two irises are attached, as far away as possible, to a 95 mm rail bracket. It is convenient to use the future holder of the scan lens (component D/E in Fig. 4). This ensures that the beam will pass through the center of the scan lens. As the mirror (component F in Fig. 4) is not installed yet, an IR detection card can conveniently be taped to the ceiling to make sure that the beam is on-axis. Of note, the resonant scanner is at its origin point when no power is being provided, but the slow galvanometer needs to be provided 0 V to position the mirror at its origin point. This can be done through the voltage supply, or to a good approximation through ScanImage by setting the scan mode to the highest possible zoom level. In our microscope design, it can be helpful to wiggle the lid to which the scanning mirrors are mounted (item C in Fig. 4). This step is tricky and can require some patience.

- When confirmed that the light travels parallel to the 95 mm rail, we attach the mirror (component F in Fig. 4), and slide it along the cage rods to a position where the beam is reflected centrally through the caging system and the mount positioned on top of the 95 mm rail. This is done using the same set of two SM2-mounted irises that we used to make sure that light leaves the scanning mirror box without x or y displacement. The beam is now ready to travel through the microscope head.

#### 2.3 Assembly of the microscope head

A picture is shown in Fig. 5.

##### 2.3.1 Components

**A)** Empty tube, shielding the beam aligned in the previous section.

**B)** Protected silver Mirror (PF30-03-P01, Thorlabs) inside KS3 and clamped down with XE25CL2 and another clamp. This mirror directs the beam through the tube lens. We will refer to this part as “big mirror”. This is the first mirror that we will have to align.

**C)** Tube lens - See detailed description below.

**D)** Empty, just a tube used for spacing because the distance between the scan lens, tube lens, and objective is important.

**E)** Tube holder (LCP01, Thorlabs) - just there to hold the lens straight, use with ER series 6 mm rods (ER6-P4, Thorlabs).

**F)** Elliptical mirror (PFE20-P01 inside KCB2EC, both Thorlabs). We will refer to this part as “collection mirror”. This is the second mirror that we will align.

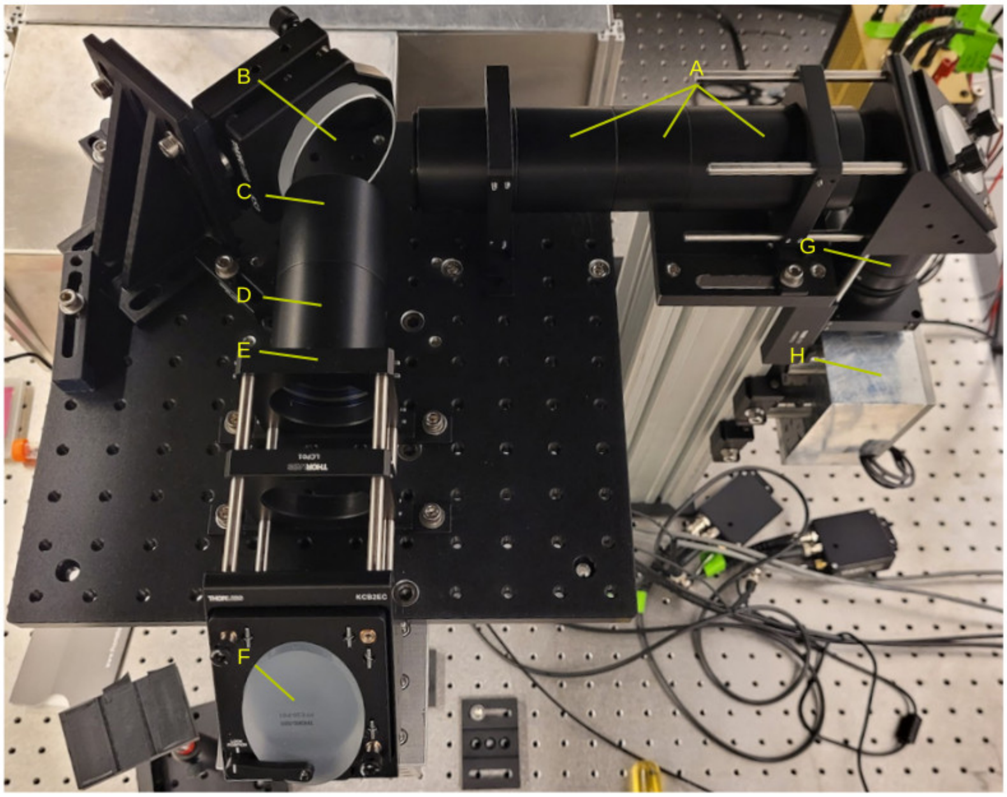

**Figure 5. Overall view of the microscope head.** Key components of the system are labeled with capital letters. These are explained in the text together with a note on how to align these components.

##### 2.3.2 Setup and alignment

- Mount the two 95 mm posts to the table.
- Mount the collection optics box to the aluminium stage, and then mount the stage to the 95 mm rail. Do not install the photomultiplier tubes yet. These devices are light sensitive, and more alignment needs to be done before we are ready to install them.
- Equip the aluminium stage with brackets and cage mounts to hold SM2 tubing (A, D and E in Fig. 5).
- First, we position the big mirror at a 45° angle (component B in Fig. 5). Place two irises after the big mirror, one at the future position of the tube lens (component C in Fig. 5) and one far away beyond the future position of the collection mirror (component F in Fig. 5). Aligning the big mirror is fiddly, but can be achieved with small adjustments of its position in x and y, and the tilt of its housing. We found it helpful to use a 45 degree angled rod to guide the mirror in the direction parallel to its surface. Whichever method is used, this step can take some time. Importantly, do not adjust the set screws of the mirror on the scanning post (component F in Fig. 4) because that is already aligned.
- Install the collection mirror (component F in Fig. 5). Then position an iris in place of the future objective, and a second iris as far away as possible just above the optical table. Use an adapter with external M32 $\times$ 0.75 threads and internal

SM1 threads (SM1A33, Thorlabs) to attach the irises to the objective collar. Then shift the collection mirror along the rail and adjust its tilt via the set screws so that the beam is aligned with the two irises.

##### 2.3.3 Scan and tube lens alignment

After the beam's alignment is confirmed by traveling through the two irises, it is time to install, and align scan and tube lens. Remember that (1) the distance between the tube lens (component C in Fig. 5) and scan lens (component G in Fig. 5) has to be 100 mm + 375 mm = 475 mm while (2) the scanning mirror set (component H in Fig. 5) has to be positioned in the focal plane of the scan lens (see Fig. 1 in the main manuscript). In our design, the objective is fixed. Therefore we have to perform the lens alignment in several steps, conceptually beginning at the back-aperture of the objective.

- Before starting, make sure the lens groups in the tube and scan lens assembly are mounted correctly. Specifically, when inserted into one of our previously aligned SM2 mounting rings, they should focus the beam, but must not incur a lateral or angular displacement. To test this, one can screw them into component E in Fig. 5, and observe the beam far away on a wall. The lens groups must not change the position of this spot, only its size. Only proceed if the lens groups were assembled and mounted correctly.

- The first lens group to be installed is the tube lens assembly. Position the tube lens such that the beam, at maximum zoom level, is focused at the back-aperture plane of the objective. As the objective is not installed yet, an IR detection card can be placed at its future location. We designed the microscope such that the tube lens assembly is to be mounted on a 1 inch long SM2 extension to the cage plates (Component D in Fig. 5. See also the diagram in the section on "Tube lens Assembly" below). Observe how the focal plane changes when moving the tube lens along the optical rails (Component E in Fig. 5). When its correct position is confirmed, tighten the set screws on the optical cage.

- Next, install the scan lens above the scanning mirror box. The spacing between tube lens and scan lens is critical because the beam leaving the tube lens has to be collimated. To test this, remove the elliptical mirror (component F in Fig. 5) from its housing, and observe the spot on a wall far away, while moving the scan lens up and down on the 95 mm optical rail. Find a position at which the beam neither converges, nor diverges after the tube lens, and the spot on the wall has the same size as the beam right after the tube lens. Only then is the beam well collimated. Clamp down the scan lens on its rail.

- Next, reinstall the elliptical mirror (component F in Fig. 5), and observe the back-aperture of the objective, while moving the scanning mirror box up and down. The scan mirrors have to be positioned in so that they are conjugated to the back aperture of the objective. It is positioned correctly when the well-collimated light converges on the back aperture of the objective across scan angles. Make sure that (1) no beam clipping is visible on high zoom along the optical path. (2) In the conjugated plane of the scan lens, a nice and crisp plane should be visible on an IR card, produced by the scanning beams focused down to a point. (3) The fast resonant scan combined with the slow scan gives the beam a visible dynamic, a "wobble". If the scanning mirror box is positioned correctly, then, the beam at the objective back aperture will not translate while scanning because all deflection angles converge on the back aperture (cf. Fig. 1 in the main manuscript). In other words, the "wobble" becomes minimal at the plane

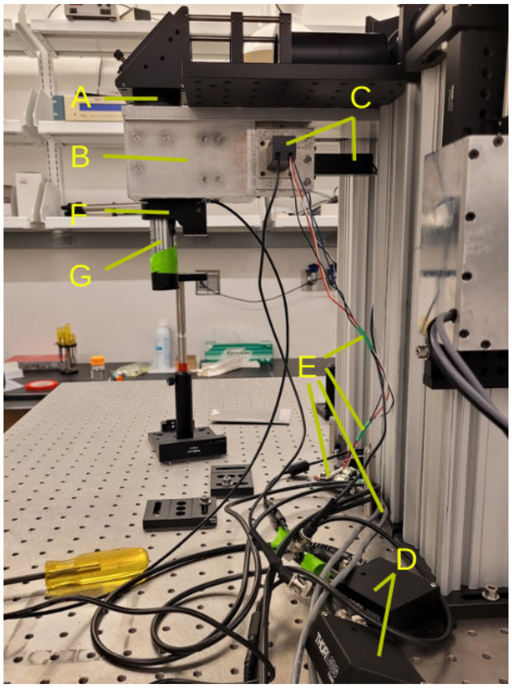

**Figure 6. Side view of the microscope head.** Key components of the system are labeled with capital letters. These are explained in the text together with a note on how to align these components.
Set up like this, we were measuring the amount of light traveling through the system. Note the detector taped to the objective with green tape.
Also note that the photomultiplier tubes (component C) are light sensitive and need to be installed in the dark. We recommend to switch off the room lights when handling them, and allowing your eyes to adapt to the dark over several minutes. After installation into the collection optics, the lights can safely be used. The objective is easy to install, and just screwed into the piezoelectric collar, component F.

conjugated to the scan mirrors. Notice how adjustment of the scan mirrors relative to the scan lens can bring this conjugate plane to any point below the collection optics. Now, adjust the height of the scanning mirror box to a point so that the minimal-wobble plane is located at the back aperture of the objective and clamp down the scanning mirror box. For a useful illustration of this method, see also [6]. Of note, this step of fine-tuning the position of the scan box up and down along the 95 mm optical rail is the reason why we stressed the importance of the alignment of the scanning post, the components A in Fig. 4 and M in Fig. 1 earlier. It was critical that across the various heights used here, the scanning mirror box was precisely aligned with the beam.

- Shield the beam with more SM1 and SM2 tubing to protect the beam from misalignment, the optics from dust, and the user from accidentally exposing themselves to the beam.

#### 2.4 Installing objective and photomultiplier tubes

##### 2.4.1 Components

The optics and optomechanics are now essentially in working order. The last missing steps towards a working microscope are the installation of the objective, and the photomultiplier tubes. These elements are visible in a side view of the microscope head, see Fig. 6.

**A)** Adjustable lens tube (SM2V05, Thorlabs), made flush with the collection box to stop room light entering the optics. Otherwise this will increase the background signal during use of the microscope.

**B)** Aluminium box housing the collection optics - See detailed description below.

**C)** Two photomultiplier tubes (“PMTs”, Hamamatsu H16201-40). Their shielded output cable (black) and the control and power supply cables (color-coded wire) are also visible.

**D)** PMT Transimpedance Amplifier (Thorlabs TIA60) - amplifier for the PMTs. Set the output offset on the PMT amplifier such that no signal equals 0 V. The amplifiers are directly connected to the PMT output on the input and ScanImage on the output.

**E)** Control signals and power supply for the PMTs going to the custom PMT control box.

**F)** Piezoelectric objective collar (P-725K085: PIFOC(R) 400 µm travel, Physik Instrumente).

**G)** The microscope's objective (Nikon N16XLWD-PF, Thorlabs).

###### 2.4.2 Objective installation

Before installing the objective, double check that the beam is collimated below the piezoelectric collar, and on-axis. To do so set the ScanImage zoom level to high and with the use of two irises confirm that the beam is not clipped. Then screw the objective into the collar.

###### 2.4.3 PMT installation

PMTs are light sensitive. Always store them with their aperture covered with blackout tape when not in use (AT205-1.0 black aluminium foil tape, Thorlabs). When removing the tape for installation, this installation into the collection box must be done in darkness and as quickly as possible. We recommend to use black silicone putty (V-tie self setting silicone rubber), and have it ready for use to seal the gap between the PMTs and the aluminium box. Switch the room lights off, wait a few minutes to adapt to the darkness then mount the PMTs on their respective ports. Of note, when ordering the PMTs from Hamamatsu, one can request PMTs with particularly low dark count and high quantum efficiency to reduce noise. We typically request a quantum efficiency of at least 45% at 520 nm, and a dark count of 4000 cps or below at 25 degree Celcius.

##### 2.5 Shield the light path

Now that your microscope is ready, finish shielding the light path with SM1 and SM2 tubing, lens tube cover (SC600, Thorlabs), and cover the optical table with sheet metal. We use bent 1 mm aluminium sheet, placed on small columns made from T-slotted rail.

#### 3 Last steps

##### 3.1 No signal?

If no picture is visible, check:

- Is a fluorescent sample under the objective? A bright simple will be useful, such as a fluorescent plastic slide (FSK5, Thorlabs). Is the sample positioned in a suitable distance from the objective? Note that the focal plane is only a few µm wide. If no fluorescence is produced in this optical slice, no signal will be visible in ScanImage.
- If the objective requires water immersion, has a drop of water been placed between the sample and the objective?

- Are all beam blocks removed from the path? Check with IR cards along the way. Below the objective, the IR light should be focused to a tight spot.
- Are the PMTs operating as intended? This can be checked with a light-source like a flashlight close to the objective. This should noticeably increase the noise in the picture.

##### 3.2 Validation and tuning

Validation and tuning is done in several steps.

- The first sample to image should be a fluorescent plastic slide (FSK5, Thorlabs). These are bright and uniform. It can be helpful to scratch the surface of the slide to find a feature to focus on. With this sample, the group delay dispersion can be adjusted for maximum signal. It should also validate that the field-of-view is as intended, and no obstructions exist in the beam path.
- Next, image a calibrated scale to validate the microscope's magnification. We used a reference slide featuring a 100 µm grid (Thorlabs R1L3S3). This slide was covered with a drop of Fluorescein solution in water and a standard coverslip. Find the calibration grid, and compute the magnification in µm/pixel. It should be approximately equal in x and y direction. Our microscope has sufficiently small field curvature that despite the thin film of dye, the entire grid should be visible. An example is shown in Fig. 7A.
- Next, measure (at very low laser power to not saturate the PMTs) the two photon intensity from a uniform bath of Fluorescein, see Fig. 7B. This measures field flatness. Alignment problems either in the excitation or collection path will make this image less uniform. Of note, small alignment errors in particular of the scan lens can have a major effect on field flatness. If your field is not as uniform as the example shown, we recommend to check the scan lens first.
- Finally, use a fluorescent bead sample to measure the microscope's point spread function (PSF), see Fig. 7C. This will reveal more subtle misalignment and aberration in the optics. We use a 5000:1 mix of 100 nm fluorescent beads ("Dragon Green" 480/520 from Bangs Laboratories) in in 1% Agarose. To prepare this sample, take the beads from the fridge, vortex the container, take the 10 µl and 49.5 g of water. Then microwave the mix with 0.5 g of Agarose until the solution is clear. While warm, place a drop on a microscope slide, and cover with a coverslip. Bring your sample to the microscope, and record a z-stack. Use low laser power to not damage the beads, or saturate the PMTs. The example shown was recorded with 5% laser power at 99.9 zoom for 40 z-slices with 0.5 µm spacing. The PSF profile should be symmetric, and a faint, but also symmetric, Airy ring visible around the bead, around 5 µm away from center of the bead (see Fig. 2D in the main manuscript).

#### 4 Assembly of subcomponents

In the remaining sections, we will cover the microscope's sub-assemblies and their construction, beginning with the lens groups.

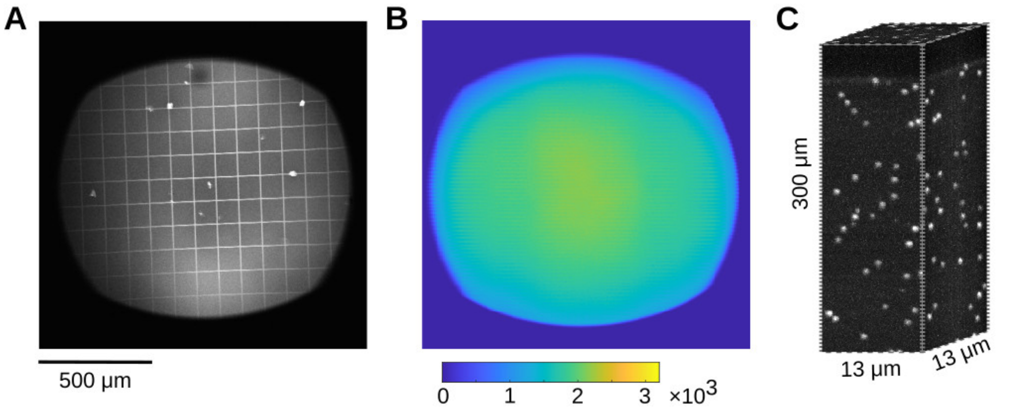

**Figure 7. Three key validation samples. A)** Image of a reference slide with a 100 µm grid to measure the microscope's magnification. Note how the entire grid is in focus, suggesting small field curvature. **B)** Two photon intensity from a uniform bath of Fluorescein. This shows that the microscope can effectively elicit and collect fluorescence across the field of view. **C)** A volumetric image of fluorescent beads in Agarose. This is used to compute the axial and radial width of the microscope's point spread function. Use low laser powers for these measurements to not saturate the PMTs and to produce correct results.

#### 4.1 Tube lens assembly

The tube lens assembly is made of two cemented doublets 2× ACT508-750-B (Thorlabs), see Fig. 8A. The letter "B" signifies that these lenses have an anti-reflective coating in the IR range. For assembly, the lenses are placed on a lens assembly tool, and a SM2 housing is lowered above them, for an example see Fig. 8B. This methods has to be used for the lenses to be seated correctly. The individual steps for the tube lens group are as follows:

- Place the first element on top of the spanner wrench (Thorlabs SPW604) using lens paper. Check the lens thoroughly. Dust can be blown away with a duster, finger prints wiped off with Methanol and a lens cleaning tissue.
- Blow it with the air duster.
- Slide the SM2L10 tube over the element until it sits in the end. Sometimes it needs to be wiggled, but do not force it.
- Flip the whole thing over.
- Now the element is in the tube. Tighten it with a SM2 retaining ring.
- Insert a second SM2 retaining ring for spacing.
- Repeat the previous steps to install the second lens above the SM2 retaining ring
- Flip again, blow with air, and set the lens in place with a third SM2 retaining ring. This produces the configuration shown in Fig. 8A.
- Be careful with dust and fingerprints when putting elements into the tube. Confirm that there are no fingerprints, and no dust particles trapped on or between the lenses.

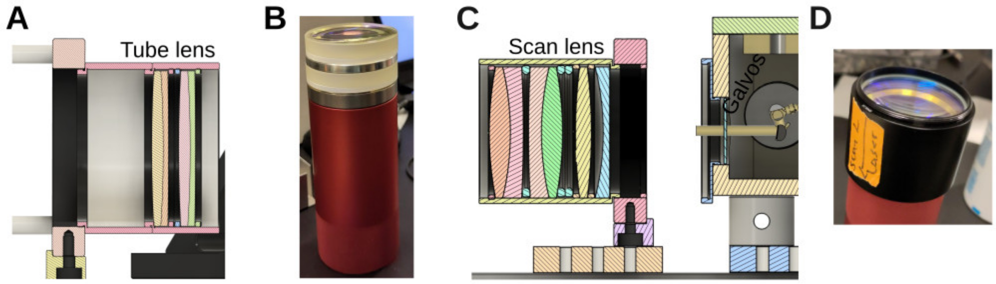

**Figure 8. Assembly of lens groups. A)** Tube lens assembly with the two doublets held in place with spacer rings. **B)** Assembly method of lenses: Lenses and spacer rings are stacked on a SM2 spanner, ready for the SM2 tubing to be lowered around them. **C)** Design of the scan lens assembly. Notice the custom spacer rings in blue. The distance between the last element of the scan lens assembly and the scanning mirror box is $\approx 1.35$ inch. **D)** The assembled scan lens group in its SM2 housing.

Our simulations indicate that the lens spacing between the two doublets is not critical, and the approximate clearance provided by two SM2 spacer rings is sufficient. This is shown in Fig. 8A. For more precise placement, a spacer ring like the one shown in Fig. 8B can be used. This will be important for the slightly more complex scan lens assembly described in the next section.

#### 4.2 Scan lens assembly

The scan lens assembly is shown in Fig. 8C. The individual steps for the scan lens group are as follows:

- Place the KPC070AR, a plano-concave lens, on the spanner wrench with the plane side facing out. Check the lens thoroughly for dust and finger prints.
- Slide the SM2 tube over the elements until they sit at the end of the tube. Sometimes it needs to be wiggled, to make sure the lenses are seated properly.
- Flip the whole thing over.
- Tighten it with a SM2 retaining ring.
- Next, place the LB1199-B lens on the wrench. This is a bi-convex lens for which the orientation is irrelevant. Flip over the assembly, and tighten the lens with a second SM2 retaining ring.
- For the third and fourth lens, place an ACT508-200-B lens, a spacer ring, the second ACT508-200-B lens, and two spacer rings on the wrench. Make sure you remove all dust from the various surfaces. Flip over the assembly, and tighten the lens with a third SM2 retaining ring.
- Check the assembly for trapped dust or fingerprints. This will produce a lens group like the one shown in Fig. 8D.

#### 4.3 Assembly of the collection optics

The collection optics are made from several components, including optical filters, lenses, and custom metal pieces. Fig. 9A shows a cut through the CAD drawing. The function is described in the main manuscript. A pictures of the collection optics box is shown in

Fig. 9B with key components labeled. Here, we will focus on the assembly and their alignment.

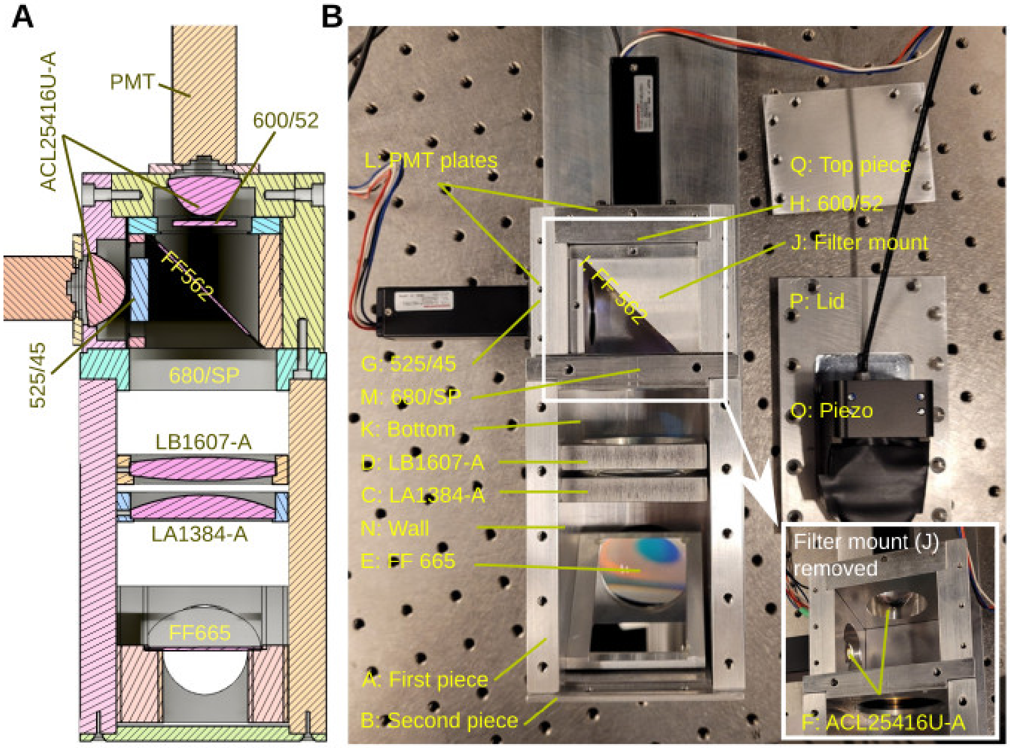

**Figure 9. Assembly of the collection optics. A)** The assembled collection optics with bottom cover removed, perspective from below. **B)** The collection optics during the assembly process, similar to panel A. Key components are labeled (see text). EDITS TO MAKE IN THE ARROWS

**A)** The construction of the collection optics box starts with a side panel attached to the base with stainless steel button head hex screws, 8-32 thread size, 1/2 inch long.

**B)** Next, we attach the front panel. This is connected with stainless steel Phillips flat head screws, 82 degree countersink angle, 4-40 thread size, 1/8 inch long and undercut.

**C)** Then we install the first lens, a LA1384-A plano-convex lens (Thorlabs), see Fig. 9. The flat side of the lens sits against the lip inside of its aluminium holder. The convex side points towards the PMTs. The lens is glued in place with three dots of Epoxy (because 3 points make a plane, and 4 can bend the lens). When gluing, make sure to dab the glue on and pull AWAY from the center of the mirror. That way any glue strands do not fall onto the mirror. When the glue has set, the holder is screwed into the wall.

**D)** The next lens to be installed is a biconvex lens (LB1607-A, Thorlabs). This lens is symmetric, so can be oriented in any way. It is also epoxied on three spots and the holder screwed into the wall, see Fig. 9A.

**E)** Next, we glue the first dichroic mirror to its holder. This dichroic mirror separates IR light from fluorescence (FF665-Di02-40X55, AVR Optics). Its orientation is important. The caret points TOWARDS the metal piece. The filter is then glued in place with three dots of Epoxy, and the triangular holder screwed into the wall.

**F)** Next, we fix the condenser lenses (ACL25416U-A, Thorlabs) in their housing plates. The convex part is facing towards the inside of the box, and set with nylon set screws (Stainless Steel Nylon-Tip Set Screw, 4-40 thread size, 1/8 inch long). The holders are then fixed together with stainless steel socket head screw, 8-32 thread size, 3/8 inch long.

**G)** With the condensers in place, we can install the optical band-pass filters in front of them. First the 525 nm band-pass filter (Semrock Brightline FF01-525/45-32, AVR Optics) for the green channel, which is secured with a nylon set screw (stainless steel nylon-tip set screw, 4-40 thread size, 1/8 inch long). Orientation is important: For this brand of filters, the arrow is following the light path, i.e. away from the inside of the box and towards the PMT.

**H)** Next we install the 600 nm band-pass filter for the red channel (Semrock Brightline FF01-600/52-32, AVR Optics). Like the 525 nm filter, it is secured with nylon set screws and the arrow points away from the inside of the box and towards the PMT.

**I)** Next, we attach the dichroic mirror that splits red and green light (Semrock Brightline FF562-Di03-40X52, AVR Optics) to its metal holder. The caret points towards the metal holder, such that the side the caret is pointing to contacts the metal when it is glued in place. When the glue has set, the mirror casing is screwed in place using stainless steel socket head screw, 4-40 thread size, 5/16 inch long. The glue used is a 5 min non-sagging Epoxy. Three dichroic was attached with 3 points of glue of minimal volume.

**J)** The piece containing the 525 nm and 600 nm band-pass filters and the dichroic mirror is connected to the larger metal box with stainless steel socket head screw, 4-40 thread size, 3/4 inch long.

**K)** The bottom piece (bottom per the view of the picture in Fig. 9B) of the PMT box is attached using 4-40 thread size, undercut, 3/16 inch long screws.

**L)** PMT plate. The PMT attaches to the flat side of the PMT plates with 4 specialized low profile screws (CBSTSR2-6, Misumi). The other side has a shallow cut out for the screw head to sit. The plate with the PMT attached screws into the rest of the assembly using 4-40 thread size, 5/16 inch long socket cap screws.

**M)** Next, we install an IR block filter (Semrock BrightLine Multiphoton Filter FF01-680/SP-50, AVR Optics). This filter is sandwiched between two rings (SM2RR).

**N)** Then we close the second panel with 8-32 thread size, 1/2 inch long socket cap screws.

**O)** The piezoelectric holder is attached to the aluminium part using the adapter that comes with it. Be careful to not damage the Serial Cable connected to the holder.

**P)** The lid is attached using 8-32 thread size, 3/8 inch long, undercut screws. Finish off by securing the rest of the front plate screws (component B) using 4-40 thread size, 1/8 inch long screws. Screw in the piece holding M to the lid using 4-40 thread size, 1/2 inch long socket cap screws.

**Q)** Attach the top cover using 4-40 thread size, undercut, 3/16 inch long screws.

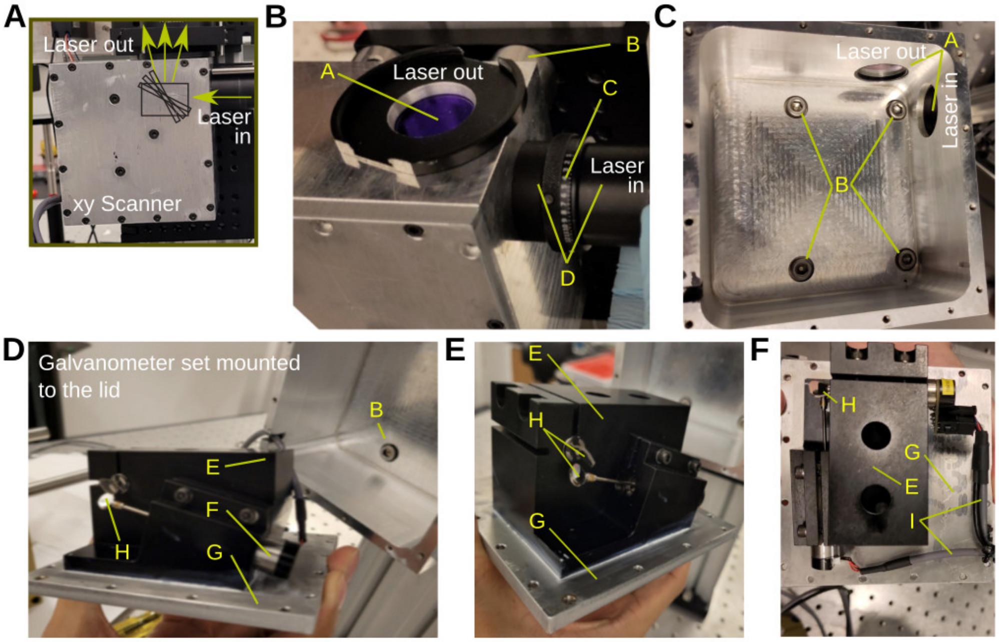

**Figure 10. The scanning mirror box. A)** The custom aluminium box housing the pair of scanning mirrors. **B)** Close-up of the same box with key parts labeled: A: Two windows (WG11010-B, Thorlabs) to keep dust out. B: Attachment of the box to the 95 mm optical rail. C: An Iris used for alignment, and D: SM1 tubing. **C)** Bottom part of the aluminium box. A: the windows, and B: screws to attach the box to the drop-on rail carriage via four 3/4 inch tall ⌀1 inch diameter posts. **D)** The scanning mirrors are mounted to the lid using heat transfer compound. E: Anodized aluminium block holding the scanning mirrors. F: One of the scanning mirror assemblies, G: The aluminium lid and H: The scanning mirrors. **E)** Same as **D** but different angle. **F)** Same as **E** but different angle. I: Cables controlling the scanning mirrors.

#### 4.4 Assembly of the scanning mirror housing

The scanning mirror housing is a machined piece of aluminium holding the Novanta scanning mirror set against the 95 mm optical rail (XT95, Thorlabs), see Fig. 10. It also shields the scanning mirrors from dust in the environment, and protects the environment from audible noise produced by the resonant scanning mirror. The housing is composed of a box, mounted to the 95 mm rail (Fig. 10A,B,C), and a lid to which the Novanta scanners are mounted (Fig. 10D,E,F). When mounting the black anodized holder, make sure to apply heat transfer compound between the holder and the lid (we use Wakefield Thermal Joint Compound Type 120-5 Silicone, cf. Fig. 10D,E). While the resonant scanner does not dissipate much heat, the slow galvanometer becomes warm in normal operation, and this heat needs to be dissipated. This is particularly important when pushing to fast responses, and driving the slow galvanometer with larger currents.

The two box apertures (beam input and output) have near-infrared antireflection-coated flat BK7 windows (WG11010-B, Thorlabs) secured with SM1 tubes and retaining rings (SM1RR, Thorlabs). We used a SM1 to SM2 converter ring at the exit, Fig. 10B, to ease checking alignment with downstream mounts and optics.

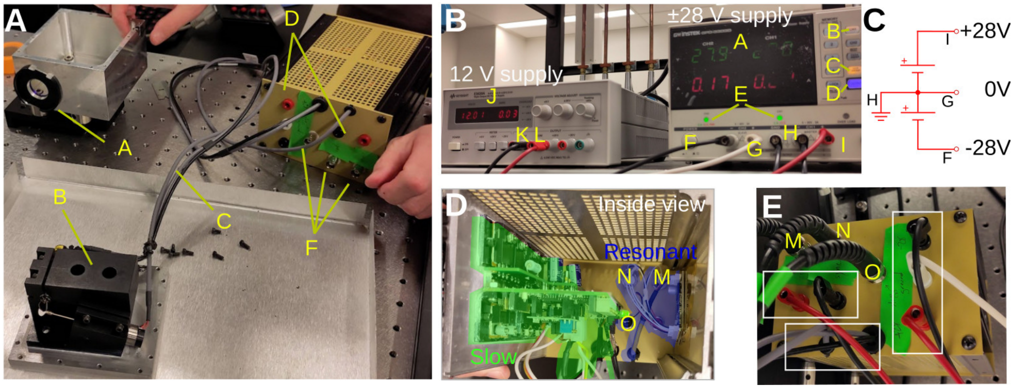

**Figure 11. Scanner electronics. A)** Overview of all scanning mirror parts. Key parts are labeled with capital letters. Optomechanics were covered in Fig. 10. All electrical drivers were placed in the yellow circuit box. **B)** Power supply and connection of the scanning mirrors with key parts labeled by capital letters. **C)** Wiring diagram for the ±28 V power supply, using two galvanically isolated supplies. **D)** The scanning mirror driver boards, provided by Novanta, and mounted in the yellow circuit box. The smaller driver circuit (blue) powers the resonant scanner. The larger circuit board (green) drives the slow galvanometer. The circuit boards are mounted to the aluminium walls with screws and heat transfer compound. **E)** Same box as in D, but from the outside and key parts labeled by capital letters.

#### 4.5 Scanning mirror power supply

The scanning mirrors come with driver circuit boards. We place them in a prototype box (EG6, Gold Box, Acopian), and connect the driver board's power supply and signal lines to connectors for easier access, see Fig. 11A. The driver boards are mounted to the aluminium walls of the prototype box with heat transfer compound (Wakefield Thermal Joint Compound Type 120-5 Silicone). The power connectors accept banana plugs. The signals are connected to BNC connectors. Fig. 11A shows as the housing box (component A) and xy scanning mirror set (component B) that we discussed above. Components C are the cables connecting the scanning mirrors to their driver boards. These driver boards are housed in a yellow prototype box, and connected to accept power (component D), and signals (component F). Fig. 11B shows the power supply of the scanning mirrors. At time of purchase, we requested the controller of the slow galvanometer mirror to be configured for usage at ±28 V (manufacturer default is ±24 V). This overall improves dynamical properties. The controller for the slow scanner requires three supplies respectively: -28 V, 0 V, and +28 V.

To provide these three signals, we used a power supply with two galvanically isolated and adjustable output channels, that can be connected in series (Note: not every power supply supports this function. We used a GPD-3303D from Gw Instek that allows this function with the SER/INDEP key). The voltage of each of the two unconnected channels was set to +28 V. To produce the three supply signals, we used these two channels in series. In other words, we connect the positive terminal of the first channel to the negative terminal of the second channel. As the channels are otherwise not connected to a common ground (this is meant with the words "galvanically isolated"), this does not produce a short circuit. Instead, this link effectively provides a new reference point, relative to which the first supply produces a negative -28 V supply, and the second supply produces the +28 V supply. This is illustrated in Fig. 11C. In this arrangement, a multimeter would read between points G and I +28 V, between G and F -28 V, and between F and I +56 V. The supply of the resonant scanners is simpler, as it only requires a single +12 V line. The supplies are connected to the power supply ports on the prototype box, in which the two driver boards reside, see Fig. 11D. Signal lines are connected on the outside with coaxial cables to the MBF Bioscience DAQ system, see Fig. 11E. Key components in the panels are labeled with capital letters. These are as follows:

- **A)** Power supply (GPD-3303D, Gw Instek) with two channels configured in series, both set to 28 V.
- **B)** Have CH 1 turned on.
- **C)** Press the "Ser/Indep" button. This wires the two channels together to a shared references to provide the ±28 V and the 0 V reference line, see Fig. 11C.
- **D)** This button enables our power supply unit. Only when pressed, power is sourced to the galvanometer driver board.
- **E)** CC/CV indicator light. Make sure that the device is set to CV (constant voltage).
- **F)** The -28 V output line (black). This is connected to the -28 V line on the electronics box (shown below in Fig. 11E).
- **G)** The reference lines, 0 V (white) connected to the 0 V line on the electronics box, see Fig. 11E.
- **H)** Connects the reference line to ground.
- **I)** The +28 V output line (red) connected to the +28 V line on the electronics box. Conform the correct voltage with a reliable multimeter BEFORE powering up the galvanometer controller, e.g. with a Fluke 117.
- **J)** Power supply (E3630A, Keysight) set to 12 V to power the resonant scanners.
- **K)** Black cable is the reference, and connected to black port on the controller box. Just below the connectors M/N/O.
- **L)** Red channel, set to +12 V, is connected to the red port on the controller box. Just below the connectors M/N/O.
- **M)** Coaxial connector controlling "Zoom" (the resonant scanners's amplitude).
- **N)** Coaxial connector for "Sync" (the resonant galvanometer's digital synchronization signal)
- **O)** Coaxial connector for controlling the slow galvanometer.

Make sure the power supply settings are correct before plugging in the scanning mirrors. Confirm this independently with a reliable multimeter (e.g. Fluke 117) and do not just trust the display. Of further note, the scanning mirror driver boards cannot be mixed and matched. Every board has a unique ID that is matched to the scanner. Double check that they are connected correctly before powering up.

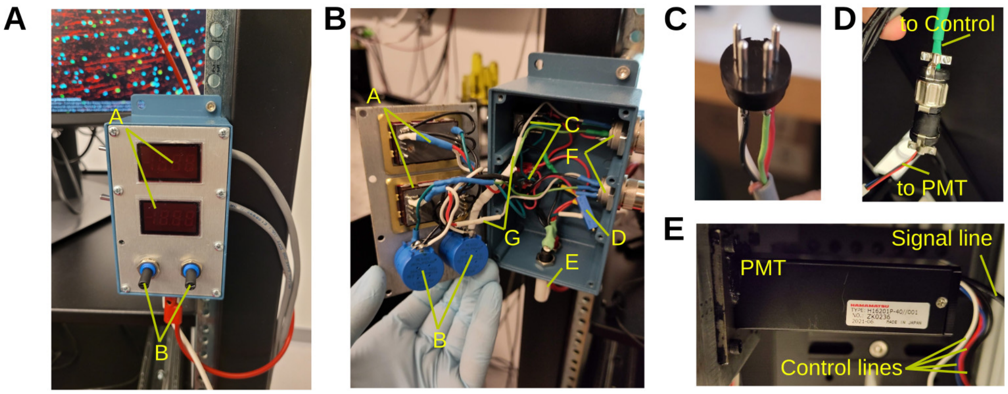

**Figure 12. Custom electronics. A)** Picture of the control box. Component A are the displays for the control voltage for red and green channels. Component B are the two potentiometers to set the two channel gains. **B)** Inside view of the potentiometer box with key components labeled (see text). **C)** Connector attached to the controller box before plugging into the PMTs. **D)** Connector from the Control box to the PMT cables. **E)** Outputs of the PMTs. Note the four color-coded American wire gauge #28 cables, and the coaxial signal line [7].

#### 4.6 Assembly of PMT control circuits

The last part to be assembled is the control circuitry for the photomultiplier tubes. This is shown in Fig. 12. The circuit is shown and explained in the main manuscript. Here, we will cover the assembly. In short, the circuit allows a user to set the gain of the H16201P-40 photomultiplier tube module with a potentiometer [7], while displaying the control voltage on a small digital panel meter (Murata DMS-20PC). The supply of this circuit is a well regulated +15 V supply that also powers the PMTs. The panel meter needs a +5 V supply that we obtain from the +15 V via a voltage divider. The PMT is supplied directly with the +15 V. It also sources a signal through the blue cable that is connected to ground via a voltage divider, using a potentiometer. This voltage divider produces the control voltage that is fed back to the module via the white input line, and used for display on the panel meter. Due to the simplicity of this circuit, we do not solder this on a circuit board, but rather in a prototype development box directly, see Fig. 12A,B, which is attached to the rig. The first components to be wired up are the Murata DMS-20PC displays. The circuit diagram is shown in the main manuscript Fig. 4.

- Solder pins 7 and 8 together on each of the two displays. The displays are component **A** in Fig. 12A,B.
- Solder 2.5 inch long and 5 inch long pieces of wires to pin 12 on the top display.
- Solder the other end of the 2.5 inch long piece of wire and a new 1 inch long piece of wire to pin 3 (+5 V return).
- Solder the other end of the 1 inch long piece of wire, and add 1.5 inch and 3 inch long pieces to display pin 6 (DP1).
- Solder the other end of the 1.5 inch long piece and another 1 inch long piece to pin 3 (+5 V return) of the bottom display.
- Solder the other end of the 1 inch long piece and a new 2 inch long piece to pin 6 (DP1) of the bottom display.

- Solder the other end of the 2 inch long piece to pin 12 (input lo) of the bottom display.
- Solder the other end of the 3 inch long piece from pin 6 (DP1) of the top display and a new 12 inch long piece of wire to the middle pin of the left potentiometer.
- Solder a 12 inch long piece of wire to pin 3 (closest to the knob) on the left potentiometer (50 kOhm, 3590S-1-503L, Bourns). These are the two big blue parts, component **B**, in Fig. 12A,B.
- Solder a 12 inch long piece of wire and 5 inch long piece of wire to pin 11 (input hi) on the top display.
- Solder the other end of the 5 inch long piece of wire to pin 1 (furthest from the knob) on the left potentiometer.
- Solder a 5 inch long piece of wire to pin 1 and pin 2 (+5 V supply) of the top display.
- Solder the other end of the 5 inch long piece of wire to the middle pin on the side opposite to where it says ON/ON on the switch.
- Solder a 12 inch long piece of wire and a 5 inch long piece of wire to pin 11 (input hi) on the bottom display.
- Solder the other side of the 5 inch long piece of wire to pin 1 (further from the knob) on the right potentiometer.
- Solder a 12 inch long piece of wire to pin 3 (closest to the knob) on the right potentiometer.
- Solder a 12 inch long piece of wire and the other end of the wire connected to pin 12 of the top display to pin 2 (middle pin) on the right potentiometer.
- Solder a 6 inch long piece of wire to the middle pin on the side adjacent to where it says ON/ON on the switch (component **C**).
- Solder a 3 inch long piece of wire to the left pin (left with the words being oriented the correct direction) on the side adjacent to where it says ON/ON on the switch.
- Solder the other end of the 3 inch long piece of wire and a 12 inch long piece of wire to the left pin (left with words being oriented the correct direction) of the second switch on the side opposite to ON/ON words.
- Solder a 12 inch long piece of wire to middle pin on the second switch on the side opposite to the ON/ON words.
- Solder a 6 inch long piece of wire to left pin on the potentiometer used for the +5 V supply of the display (component **D**).
- Solder a 6 inch long piece of wire to right pin on the same potentiometer.
- Solder a 4 inch long piece of wire to center pin on the same potentiometer.
- Connect the left and right pins to the banana plugs and set the potentiometer to 5 V going across BEFORE connecting the middle wire to the rest of the controller circuit!
- Solder the other end of the 4 inch long wire and another 4 inch long piece of wire to the middle pin on the second switch on the side adjacent to the ON/ON words.

- Solder the other end of the 4 inch long piece of wire to the right pin (right with words being oriented in the correct direction) on the side opposite to the ON/ON words on the first switch.
- Solder a 5 inch long piece of wire to the right pin (right with words being oriented the correct direction) on the side adjacent to where it says ON/ON on the second switch.
- Solder the other end of the wire to pin 1 and pin 2 (+5 V supply) on the bottom display.
- Trim the wires from the middle pin of the gain adjusters by around 5 inch, connect those two wires, the wire furthest from the dial on the potentiometer, and two 12 inch long wires by twisting together.
- Solder a 3 inch long wire to the white banana plug and screw it in the housing. This is component **E** in in Fig. 12B.
- Solder a 3 inch long wire to the red banana plug and screw it in the housing.
- Connect the twisted together wires to the white banana plug.
- Connect the wire closest to the dial on the potentiometer and the left pin (left with words being oriented in the correct direction) on the second switch on the side opposite to the ON/ON words to the red banana plug.
- Solder one of the wires from the white banana plug to pin 2 of the top 4-pin female connector. This connection is item **F** in Fig. 12B. The 4-pin connector that will plug into this port is shown in in Fig. 12C.
- Solder the wire from the pin closest to the knob on the left potentiometer to pin 3 of the top 4-pin female connector. Put a 12k Ohm resistor within the path of the wire (component **G** in white heat-shrink tubing). This limits the control voltage to approx. 0.8 V, which is safer for the PMT [7]. We discuss this in the main manuscript.
- Solder the wire from pin 11 on the top display to pin 1 of the top 4-pin female connector (M16 4 Pin Connector).
- Solder the wire from the middle pin on side adjacent to ON/ON words on the top switch to pin 4 of the top 4-pin female connector.
- Do the previous 4 steps for the other 4-pin female connector.

This concludes the assembly of the control box. The connection of this box to the PMTs is made through a custom cable. A male connector, shown in in Fig. 12C, plugs into the control box on one end, and into a female plug soldered to the PMT module on the other. This is shown in in Fig. 12D. The PMT module has four colored control lines attached to, see in Fig. 12E, that are wired to this connector. The specific type, shown in Fig. 12D, is a Wire-Pro, Inc. (WPI) 126-195 connector.

### 5 Software and control

#### 5.1 Laser settings

The laser control depends on the system used. The Alcor 920 laser is controlled through a USB interface, and software that runs on the same PC as ScanImage. The software is used to switch on the laser, switch between alignment and full-power mode, and to adjust the group delay dispersion.

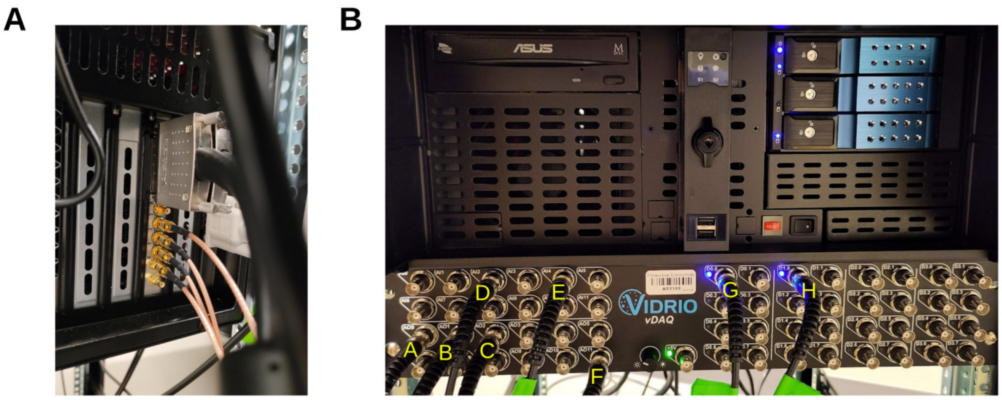

**Figure 13. Setup Vidrio DAQ system. A)** Wiring of the four input channels. These are connected to the PMT preamplifiers, and for our system digitized at 14 bit resolution. **B)** Data acquisition panel connected to the microscope's various control and signal lines. Capital letters are explained in the text.

#### 5.2 Shutter settings

The shutter controller has various settings that allow either manual control, or control via ScanImage. In our system, a "Local/Remote" toggle switch selects between those options. A second important toggle switch is "N.O./N.C.". In the "N.C." position, the shutter will be activated open by an input pulse signal. In the "N.O." position the shutter will be activated closed. If ScanImage's shutter control signal is not correctly actuating the shutter, very likely this toggle switch needs to be flipped.

#### 5.3 Setting up ScanImage

At this point, all hardware is set up and aligned. The next step is configuring ScanImage, the software used to control the microscope [4, 5] to communicate with the various components. This is done through a number of coaxial lines that connect the Vidrio data acquisition system with the microscope hardware, see Fig. 13. Our system is connected, on the back, to the two PMT outputs, see Fig. 13A. The front panel links the system to eight signals, Fig. 13B. These are

- **A)** `Analog-out`: X-resonant zoom control.
- **B)** `Analog-out`: Y-slow control.
- **C)** `Analog-out`: Piezo control for the objective collar.
- **D)** `Analog-in`: Piezo feedback from the objective collar.
- **E)** `Analog-in`: Pockels cell feedback.
- **F)** `Analog-out`: Pockels cell control.
- **G)** `Digital`: Resonant scanner sync.
- **H)** `Digital`: Shutter control.

When connected, ScanImage needs to be configure to make use of these connections. The process is as follows.

- Type `ScanImage` into Matlab and press Enter.

- Login using the correct credentials.
- Under `Machine Data File`, click `New`.
- Type in a filename and click `Save`.
- Select `Blank Configuration`.
- Next we need to add all the devices of the microscope. Start by clicking the `Devices` tab, and pressing `+` to add each new device.
- Shutter
  - Select `Digital Shutter` and click `Add`.
  - Type in a name like `Shutter1`.
  - In the new window, select `/vDAQ0/D1.0` for `Control Channel`. This will depend on how you have wired up the control line to the front panel.
  - `Open time (seconds)` remains at the default of 0.5 s.
  - Check the `Invert Output` box.
  - Click `Apply`.
  - Make sure the shutter itself (on the driver box) is on `STD` for the key and the toggle switches (see above) are set to `N.C.` and `Remote`.
- Pockels cell:
  - Select `Pockels Cell` and click `Add`.
  - Type in a name like `Pockels1`.
  - In the new window, select `/vDAQ0/AO11` for `Control Channel`. Again, this will depend on the channel that the Pockels driver is connected to.
  - For `Feedback Channel (optional)`, select `/vDAQ0/AI4`. This is connected to the photodiode directed at the beam block for the shutter. The gain should be chosen for a good signal. In our system, a gain of 50 dB works well. If the ambient room light is too bright, a SM1-mounted infrared long-pass filter in front of the photodiode can help (e.g. FELH0850, Thorlabs). The goal of setting the gain is sigmoidal control curve in ScanImage: Light transmission starts at a minimum, and increases as the control voltage increases.
  - Under `Control output [Volt]`, change `Max` to `1.7`. This is the maximum control voltage applied to the Pockels cell driver.
  - When calibrating, notice how it takes about 1.7 V to reach 100% laser power.
  - Leave the `Open shutters for calibration` field unchecked because our shutter is positioned after the Pockels cell and the reflection off the closed shutter used for calibration with a photodiode.
  - Leave everything else in the `Basic` tab unchanged.
  - Leave everything else in the `Calibration Settings` and `Advanced` tabs unchanged.
  - Click `Apply`.
- Slow galvanometer:
  - Select `Analog Galvo` and click `Add`

- Type in a name like `Y-Galvo`
- In the new window, select `/vDAQ0/AO1` for `Control Channel`. This will depend on the port that the galvanometer is connected to.
- The rest of the defaults have to be adjusted as per the Galvanometer's specifications. For our slow galvanometer, these are as follows.
- Change `Volts per optical degree [V/deg]` to `0.7692`.
- Change `Angular range [optical degrees]` to `26`.
- Change `Park position [optical degrees` to `13`.

• Resonant scanner:

- Select `Resonant Scanner` and click `Add`.
- Type in a name like `X-res`.
- In the new window select `/vDAQ0/D0.0` for `Sync Channel`. This will depend on your wiring.
- For `Zoom Control Channel`, select `/vDAQ0/AO0`.
- Leave everything else unchanged. We used a 8 kHz scanner.
- Click `Apply`.

• Piezoelectric objective collar.

- Select `PI E709` and click `Add`.
- For `Serial port / USB`, select `COM1`.
- For `Control Channel`, select `/vDAQ0/AO2`. This will depend on your wiring.
- For `Feedback Channel`, select `/vDAQ0/AI2`. This will depend on your wiring.
- Click `Read From Device` and numbers should automatically update.
- Click `Apply`.
- Click `Calib` on the new Piezo widget.
- Should be able to feel piezoelectric mount vibrate with hand when it calibrates.
- Check `Enable Stack` on the main ScanImage window and the `STACK CONTROLS` windows should pop up.
- Better to do `step` instead of `sawtooth` for resonant scanner since we are averaging images; with `sawtooth`, we would be averaging difference Z depths together
- Select 10 `Frames per Slice`, 5 `Number of Slices`, 10 `Number of Volumes`, and 5 for `Step Size [um]`.
- Click `Test Waveform`.
- In new window, click `Optimize Waveform`.
- Ringing at the beginning and end of steps can be corrected with `Actuator lag`.

• After adding the devices, we have to create an imaging system by clicking the `ScanImage` tab and click `+ Add Imaging System +`.

• In the new window, under `vDAQ Scan System`, click `Add` and type in a name like `My microscope`.

- In the new window, select `X-res` under `Resonant Scanner`.
- For `Y-Galvo`, select `Y-Galvo`.
- Check off `Use` for `Shutter1`.
- Check off `Use` for `Pockels1`.
- Check off `Use` for `Piezo1`.
- Under `Channel`, check off `Invert` for all 4 channels.
- Leave everything under the `Triggers`, `Advanced`, and `I2C` tabs unchanged. Of note: The setting `I2C` is used for syncing with behavior.
- Click `Apply`.

### 6 Recommendations for use

Make a point of regularly checking your microscope. Lenses can get dirty, the path misaligned, and components age.

- Measure (1) the laser's power levels, (2) the Pockels cell calibration and (3) check exposed lenses and mirrors for dust and fingerprints with a flashlight. This is fast and should be done before every experiment.
- When the microscope is not in use, it is good practice to switch off all components, except the scanning mirror power supplies, the PMT preamplifiers, and the photodiode.
- Every once in a while, switch the red and green PMTs to check for signs of aging and validate their performance on a fluorescent slide or sample of known brightness. The PMTs can degrade and may need to be replaced, depending on use, after several years. They should be treated with care: power ramped up and down slowly, and saturation should be avoided. We use a rotation system in which new PMTs are typically used for the green channel. When their performance becomes worse, they often have a few more months or years in the (typically brighter) red channel in them.
- It is recommended to periodically image a bead sample to validate the PSF, as well as a Fluorescein bath to detect possible misalignment. For a detailed protocol, see Lees et al. [8].
- Be mindful of imaging artifacts. When suspecting an issue, go back to the basics, and isolate the error.

For safety, it is good practice to shield the laser path. In addition to safety concerns, it will also help in keeping dust and dirt away from optical components.

### 7 Deriving the relationship between intensity and dispersion

We stated that the expected fluorescence $I(\varphi)$ as function of group delay dispersion $\varphi$ applied to a $\varphi_0$-dispersed Gaussian pulse of full-width-at-half-maximum $\Delta\tau$ can be written as

$$I(\varphi) = I_0 \left(1 + 16\log(2)^2 \frac{(\varphi+\varphi_0)^2}{\Delta\tau^4}\right)^{-1/2}. \tag{1}$$

To derive this expression, consider the electrical field of a Gaussian pulse of amplitude $E_0$, width $\Delta t$, center frequency $\omega_0$, phase $\Theta$, and centered at time $t = 0$:

$$E(t) = E_0 \exp\left(-\frac{t^2}{4\Delta t^2}\right) \cdot \exp\left(-i\left(\omega_0 t + \Theta\right)\right). \tag{2}$$

Its intensity is

$$P(t) = |E(t)|^2 = |E_0|^2 \exp\left(-\frac{t^2}{2\Delta t^2}\right). \tag{3}$$

In the Fourier domain, the spectral content of this wave package is:

$$\tilde{E}(\omega) = E_0 \cdot \sqrt{2}\Delta t \, \exp\left(-i\,\Theta - \Delta t^2(\omega-\omega_0)^2\right). \tag{4}$$

In the linear regime, frequency-dependent dispersion changes the electric field to

$$\tilde{E}(\omega) \to \tilde{E}(\omega) \cdot \exp\left(i\,\phi(\omega-\omega_0)\right) \tag{5}$$

with phase shift $\phi(\omega-\omega_0)$. We approximate this function to quadratic order $\phi(\omega-\omega_0) \approx a + b(\omega-\omega_0) + \frac{c}{2}(\omega-\omega_0)^2$. With this approximation, we obtain the Fourier representation of the electric field of the dispersed pulse:

$$\tilde{E}^{\mathrm{disp}}(\omega) = \tilde{E}(\omega) \cdot \exp\left(i\left(a + b(\omega-\omega_0) + \frac{c}{2}(\omega-\omega_0)^2\right)\right). \tag{6}$$

Coefficient $a$ causes a phase shift and coefficient $b$ a time shift. Both do not change pulse width. Computing the inverse Fourier transform to obtain $E^{\mathrm{disp}}(t)$ allows to compute the intensity of the dispersed pulse as

$$P^{\mathrm{disp}}(t) = |E^{\mathrm{disp}}(t)|^2 = |E_0|^2 \frac{2\Delta t^2}{\sqrt{c^2+4\Delta t^4}} \exp\left(-\frac{2(b+t)^2\Delta t^2}{c^2+4\Delta t^4}\right). \tag{7}$$

Comparison with Eq. (3) then shows that the pulse has broadened in time to $\Delta t \to \frac{\sqrt{c^2+4\Delta t^4}}{2\Delta t}$. Rewriting the widths of the dispersed pulse in the more common full-width-at-half-maximum $\Delta\tau$, using $\Delta t \to \Delta\tau/\sqrt{8\log(2)}$ for a Gaussian pulse, we obtain

$$\Delta\tau^{\mathrm{disp}} = \frac{\sqrt{\Delta\tau^4 + 16\log(2)^2 c^2}}{2\Delta\tau\sqrt{\log(4)}}. \tag{8}$$

The two-photon fluorescence intensity $I$ is proportional to the inverse of the pulse width, and the group delay dispersion is composed of the original dispersion $\varphi_0$ and the applied correction $\varphi$, so that $c \to \varphi_0 + \varphi$. We obtain

$$I(\varphi) \propto \frac{1}{\Delta\tau^{\mathrm{disp}}} = \frac{2\Delta\tau\sqrt{\log(4)}}{\sqrt{\Delta\tau^4 + 16\log(2)^2(\varphi_0+\varphi)^2}} = \frac{2\sqrt{\log(4)}}{\Delta\tau} \frac{1}{\sqrt{1+16\log(2)^2\frac{(\varphi_0+\varphi)^2}{\Delta\tau^4}}}. \tag{9}$$

Absorbing the constants $\frac{2\sqrt{\log(4)}}{\Delta\tau} \to I_0$ yields Eq. (1). The same result for pulse broadening, with $16\log(2)^2 \approx 7.68\ldots$, is stated as Eq. (3) in [9].
